# *CovSite*: A High-Throughput Blind Covalent Screening Framework for Reactive Site Detection

**DOI:** 10.64898/2026.07.17.739288

**Authors:** Andrew Hu, Joseph S. Bailey, Søren C. Spina, Gowrish Rajagopal, Nathan Phan, Blaise R. Kimmel

## Abstract

Targeted covalent inhibitors are a powerful, yet underexplored, class of therapeutics, and current computational covalent screeners are constrained in early drug discovery due to the need for prior knowledge of the target site and limited throughput. We present *CovSite*, a blind covalent screening tool that identifies candidate reactive residues across the entire protein surface, utilizing only the protein structure and electrophile SMILES. *CovSite* applies a pipeline of four orthogonal physicochemical filters (nucleophile identification, solvent accessibility, environment-dependent deprotonation prediction, and semi-quantum-mechanical reactivity ranking) to identify potential small-molecule candidate inhibitors. Validated against 2,062 diverse covalent protein-ligand complexes spanning six nucleophilic residue types, *CovSite* achieves a 98.5% blind target site hit on a held-out benchmark set of 207 cysteine-targeted complexes while reducing the search space by 97.8%. The target-site hit detection exceeds the 53-62% accuracy of popular covalent screening tools operating under non-blind conditions on the same benchmark set. By extending nucleophilic coverage beyond cysteine to include serine, threonine, lysine, histidine, and tyrosine, and completing a screening of a 200-residue protein in two to three minutes on standard hardware, *CovSite* serves as a platform technology with the potential to address critical gaps in throughput, generalizability, and accuracy in this field of covalent screening. We demonstrate this capability by using *CovSite* as a blind, ligand-specific approach that enables iterative, machine-learning-driven covalent inhibitor generation that is impractical with existing tools, establishing a foundation for computationally guided covalent drug discovery for novel and understudied targets.

**TOC Figure:** 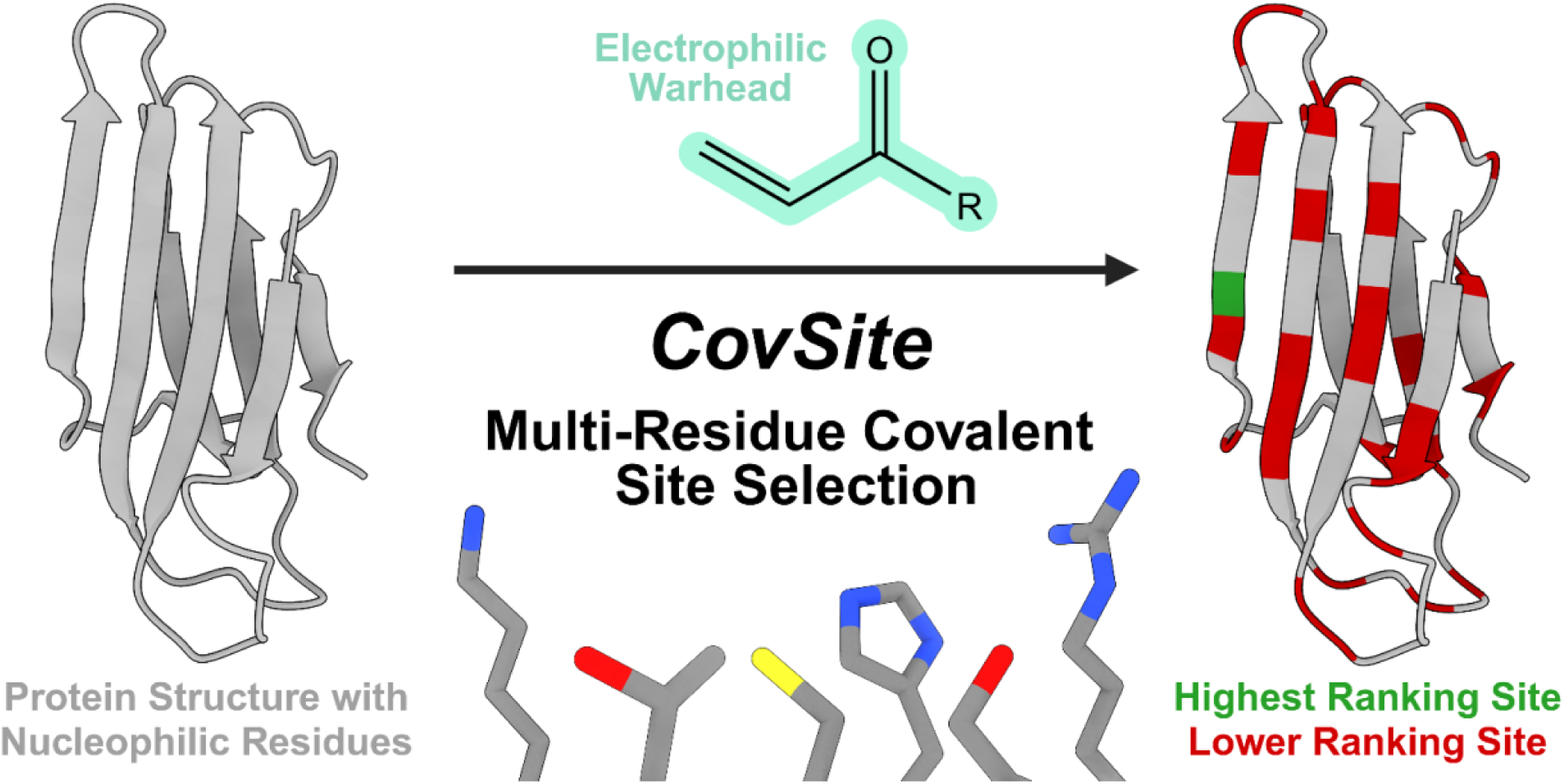

## Introduction

Recent advances in structural biology and computational modeling have enabled the rational design of targeted covalent inhibitors (TCI) for protein drug discovery. Once avoided due to concerns about toxicity, immune sensitization, and off-target effects^1^, covalent inhibitors have become powerful tools for the discovery and optimization of targeted therapeutics over the past decades. Unlike reversible inhibitors, TCIs offer prolonged target engagement, increased binding affinity, and access to previously “undruggable” protein classes^2–4^. Despite these advances, covalent drug development remains challenging. Well-studied TCIs, such as afatinib, require costly and time-consuming experimental screens to balance reactivity, selectivity, and pharmacokinetic properties. From 2020 to 2024, only 6 of 127 FDA-approved small-molecule drugs were targeted covalent inhibitors. High-throughput experimental screening of compound libraries is infeasible, underscoring the need for reliable, efficient computational tools to improve prediction accuracy^5^.

Covalent virtual screening is dominated by structure-based docking tools developed as extensions of conventional noncovalent architectures (AutoDock, GOLD, ICM-Pro, Glide)^5,6^. Conventional screening targets one of a handful of known or predictable binding pockets, as proteins typically present only 3-5 pockets large enough for small-molecule engagement that are evaluable at the pocket level without resolving environmental effects on a single residue ^6,7^. Covalent screening operates differently. Each nucleophilic residue must be evaluated independently, and chemically identical residues (e.g., two cysteines) can differ in reactivity due to local environment, a distinction that pocket-level analysis cannot capture. Because docking-based screeners are built to predict bound structures and are noncovalent docking frameworks at their core, the ligand is typically docked noncovalently, and warhead-nucleophile distance becomes the primary metric for bond formation^5,8^. Covalent bond reactivity is not incorporated into scoring^5,8,9^, and bond distance indicates geometric feasibility, not reaction favorability. Existing frameworks are not designed to predict intrinsic covalent reactivity. Reaction favorability requires quantum-mechanical (QM) calculations, which are orders of magnitude more expensive than the intermolecular potential used in docking, making explicit covalent scoring a bottleneck for throughput^5,8,9^.

These docking-based tools remain essential for resolving structures of bond formation once a reactive site is known or suspected; however, their noncovalent docking architecture makes this “known-pocket” constraint inherent rather than incidental. Several bottlenecks, including requiring prior specification of the reactive site, omitting covalent bond formation from scoring entirely, and avoiding QM calculations to preserve throughput, together mean these tools validate the geometric plausibility of a proposed covalent pose rather than the chemical favorability of covalent bond formation itself, leaving docking-based screeners prone to false positives when distance is mistaken for reactivity. The development of a tool capable of scanning a full protein surface for the reactive site is therefore of high interest, particularly for novel targets where the reactive residue is unknown, selectivity is of concern, or cryptic pockets previously considered undruggable may be involved^8,10^. This leaves a complementary gap for non-structure-based screening tools purpose-built for the covalent screening problem: determining how favorable a residue’s reactivity is, finding specific sites without prior knowledge, and narrowing the search space to reactive sites so that the remaining sites can be tested for selectivity through downstream covalent docking or experimental follow-up. A reactivity-ranking approach is positioned to close this gap directly, since scoring intrinsic chemical favorability rather than pocket geometry circumvents each of the constraints above. The reactive site does not need to be specified in advance, no QM-driven throughput penalty is incurred at the pose-scoring stage, and the output is a ranked, blind assessment of which residues are reactive, a determination that current structural screeners are not built to make.

Here, we present *CovSite*: a computationally lightweight, fully blind covalent reactive-site screening framework that requires no prior knowledge of the binding site, reactive residue, or warhead identity. Rather than adapting noncovalent docking infrastructure, *CovSite* applies physicochemical features of the ligand’s warhead, including covalent bond reactivity tailored to the covalent screening problem, to rank candidates across the full protein surface. *CovSite* integrates solvent-accessible surface area and desolvation cost, nucleophile protonation state, and intrinsic electrophile/nucleophile reactivity through Fukui indices, hard-soft acid-base (HSAB) theory, and frontier molecular orbital (FMO) descriptors to rank reactivity rather than relying on geometric proximity, thereby refining the search space and surfacing reactivity hotspots for empirical study. By identifying chemically compatible residue-warhead pairs, these results support a common covalent drug discovery strategy of attaching a viable warhead to a known non-covalent binder. Validated against 207 structurally diverse covalent protein-ligand complexes, *CovSite* correctly identified the experimentally covalent binding site in 98.5% of cases while reducing the amino acid search space of each complex by over 98%, with the true site median ranked 2nd among candidate residues. Existing tools report top-ranked pose accuracy in hit detection of 40-60% on the same 207-complex benchmark, based on previous literature, using warhead-residue distance, the field’s standard for site geometric viability for a covalent bond. *CovSite* substantially outperforms it despite operating blind. The high-throughput framework requires a minimum of only 2 user-specified inputs, the electrophile and protein, and completes screening of a 200-residue protein in 1-2 minutes on standard hardware. To our knowledge, this is one of the first covalent screening tools capable of blind reactive-site prediction with this accuracy, addressing a high-priority gap in the field. The speed, blind search capability, and accuracy of *CovSite* position it as a validation tool for generative covalent drug design pipelines, enabling screening of novel electrophilic candidates for a target protein, previously infeasible given the throughput, cost, and prior knowledge required by existing screeners.

## Results

### CovSite: A Learned Ranking Framework for Covalent Reactive Site Prediction

Covalent inhibition follows a two-step process involving a bond-forming functional group, also known as the electrophilic “warhead,” and a nucleophilic amino residue for target recognition and selective binding. First, a non-covalent bond between the inhibitor and the enzyme positions the “warhead” near a reactive nucleophilic amino acid residue (most often cysteine); then, a chemical reaction between the nucleophile and the electrophile forms residue-specific, proximity-based bonds. *CovSite* is a fully blind, ligand-aware computational framework for identifying candidate covalent reactive sites across a protein surface. Given only a protein structure (PDB file) and an electrophile SMILES string, *CovSite* computes a physicochemical feature vector for six common nucleophile-capable residues, including Cys, Ser, Thr, Tyr, Lys, His, and uses a learned ranking model to score and rank these residues by their likelihood of being the true covalent target^10^ (**Figure 1**).

**Figure 1.**
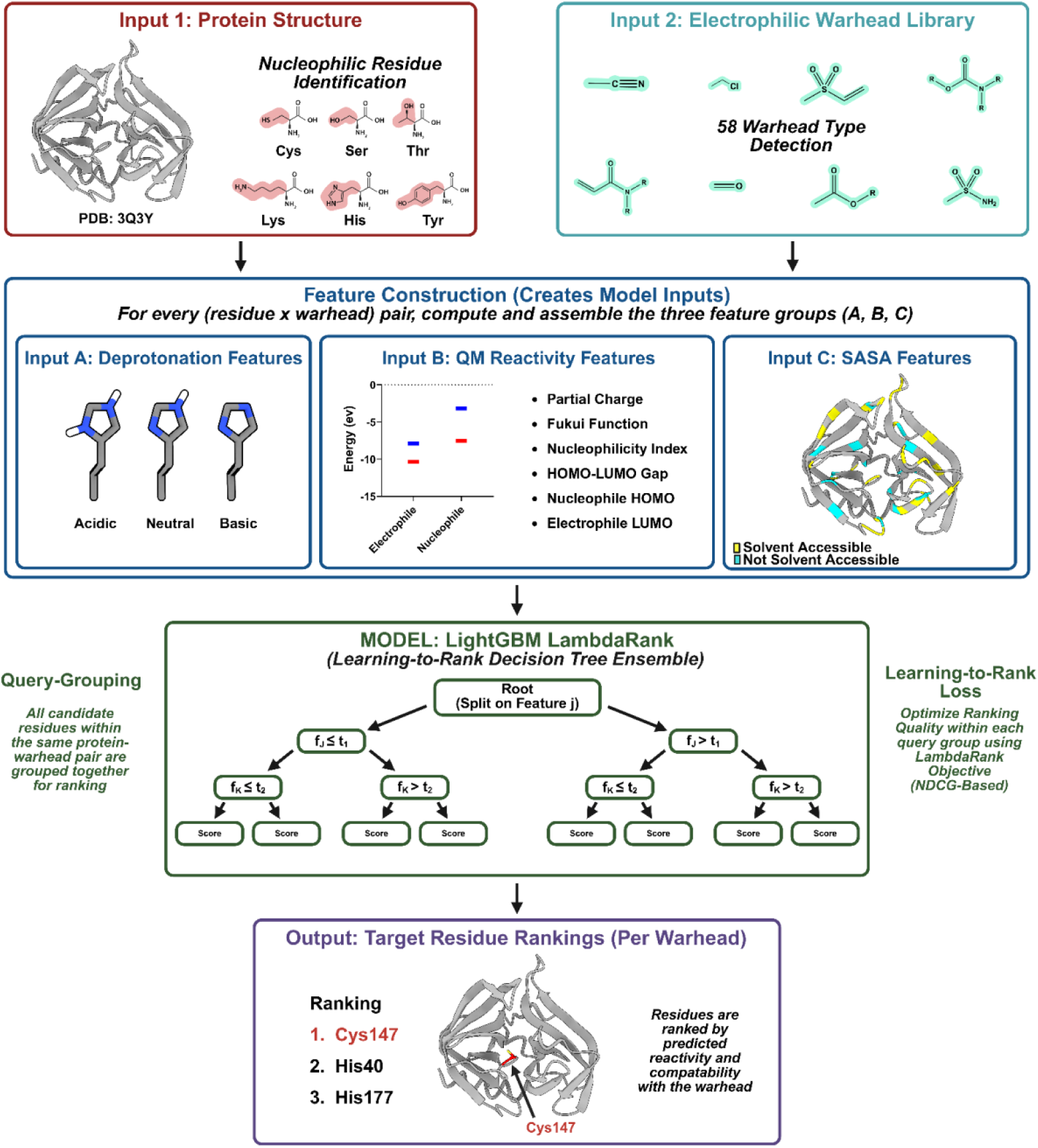
Schematic of the *CovSite* framework, including both inputs (a PDB file and ligand SMILES), the features utilized to identify the target residues on the protein, and an output candidate rank list of optimal residue positions for covalent bonding.

### CovalentInDB 2.0 Dataset Curation and Benchmarking

The *CovSite* framework was developed and tested against covalent protein-ligand crystal structures from the *CovalentInDB 2.0* database^2^. The initial dataset comprised 3,598 entries, each pairing a protein structure with an electrophile SMILES and an experimentally confirmed covalent target residue. Entries sharing a protein identifier with the independent benchmark dataset (see Independent Benchmark Evaluation below) were removed to prevent data leakage, reducing the dataset to 2,143 entries. Entries were further restricted to those targeting the six canonical nucleophilic residues of Cys, Ser, His, Thr, Tyr, and Lys^10^. Further entries were excluded when the annotated warhead chemotype could not be matched by the supported SMARTS class, commonly because the dataset’s ligand SMILES represented post-reaction adducts or prodrug/inactive drug forms, such as bisulfite adduct prodrug precursors^12^. 623 entries with inactive ligand forms were corrected to their reactive form, yielding a final curated set of 2,062 entries (**Supplementary Figure 1**).

To evaluate *CovSite* against other covalent screening tools on a common reference, hit-detection performance was additionally assessed on the C207 set. This is a set of 207 cysteine covalently targeted complexes curated by Scarpino et al. as a benchmark for comparing common covalent screening programs. Where C207 entries contained post-reaction adduct SMILES (since many covalent docking tools require a post-reaction form to specify the covalent bond), ligands were converted to their reactive electrophile form. All proteins in the C207 set (and the two case studies in the Proof-of-Concept section below) that overlapped with the *CovalentInDB 2.0* dataset were removed during curation, leading to an independent, held-out assessment. For each complex in the dataset, *CovSite* was run in blind mode, with no prior information or specification of binding site, residue, or warhead identity, and performance was evaluated by whether the experimentally confirmed covalent residue appeared in the final candidate set. Search space reduction is also calculated to determine selectivity.

### Feature Generation

For every nucleophile-capable residue (Cys, Ser, Thr, Tyr, Lys, His) on a protein surface, *CovSite* computes features spanning four physicochemical phenomena. *Solvent accessibility* is captured as the relative side-chain solvent-accessible surface area, computed with *FreeSASA* as the ratio of each residue’s side-chain SASA to its reference maximum. The *deprotonation state* is represented by the probability that a residue’s nucleophilic side chain occupies its deprotonated, reactivity-competent state, as predicted by an in-house *XGBoost* classifier described below. *Quantum reactivity* is described by five semi-empirical quantum mechanical (SQM) descriptors computed with xTB (GFN2-xTB, GBSA water solvation) for each candidate residue in its deprotonated form against the warhead identified from the input SMILES: HSAB-derived nucleophilicity index, Fukui electrophilicity index, HOMO–LUMO gap, partial atomic charge on the nucleophilic atom, and electrophile LUMO energy. An additional descriptor, nucleophile HOMO energy in the deprotonated state, is included as a sixth QM feature. *N-terminus status* is encoded as a binary indicator flagging whether a residue occupies the protein or chain N-terminus, included because the free α-amino group at certain N-termini (e.g., Thr1 in Ntn-hydrolases and proteasome subunits) is itself a reactive nucleophile independent of side-chain chemistry, and is not captured by side-chain-centric descriptors. Residue identity is additionally encoded as a one-hot vector (Cys, Ser, Thr, Tyr, His, Lys) so the model can learn residue-type-specific reactivity baselines jointly with the continuous descriptors above.

### Deprotonation Model

Because crystal structures do not resolve hydrogen positions and local electrostatic microenvironments shift effective pKa substantially from solution reference values, we trained a dedicated classifier to estimate each candidate residue’s probability of occupying its deprotonated, nucleophile-active state, rather than relying on conventional pKa prediction tools. Furthermore, the limited accuracy of current pKa tools has hampered target-site evaluation of covalent bonds, supporting the development of our own deprotonation model specific to nucleophilic residues^13^.

An *XGBoost* classifier was trained on the PKAD-R experimental pKa database with 10-fold cross-validation, using features including reference pKa for the residue type, *FreeSASA*-derived side-chain relative solvent accessibility, counts of charged neighboring residues within a set spherical radius, and hydrogen-bond donor/acceptor scores derived from local structure (weighted and strict/flexible H-bond counts). Benchmarked against PROPKA on the PKAD-R dataset, PROPKA failed to predict environment-driven protonation state changes for any of the six canonical nucleophilic residue types, performing at the level of the reference pKa baseline alone. The trained deprotonation classifier retained meaningful sensitivity to such environment-driven shifts (**Supplementary Figure 2**), by estimating a back-calculated pKa, pKₐ(est), using the following equation: pKₐ(est) = pH − log[P / (1 − P)], which is passed directly into the ranking model.

### Ranking Model

Residue-level features were combined using an *LGBMRanker* (LightGBM, lambdarank objective) trained to rank all nucleophile-capable residues within a protein–ligand complex by their likelihood of being the experimentally confirmed covalent target. Each protein–warhead pair constitutes one ranking query group; all candidate residues within that complex are scored relative to one another, consistent with the framing that covalent reactivity is only meaningfully compared among nucleophiles co-present in the same structure. Sample weights were applied per query group according to the rarity of the labeled target residue type in the training distribution, upweighting underrepresented residue types (Tyr, Thr, His, Lys) relative to the dominant Cys and Ser classes to prevent the model from defaulting to a residue-frequency prior.

### Train/Test Splitting and Cross-Validation

To prevent data leakage from homologous proteins appearing in both training and evaluation sets, protein sequences were clustered using *MMseqs2* at a 50% sequence identity threshold (80% alignment coverage) for any chains when comparing structures, a conservative approach that accounts for multi-subunit complexes. A 20% held-out test set was assigned at the cluster level (not the individual-entry level), ensuring that no training protein shares ≥50% sequence identity with any chain of any test-set protein. The remaining 80% of clusters were used for model development with 5-fold *GroupKFold* cross-validation, again grouped by cluster identity, so that every fold’s validation split is drawn from clusters entirely unseen during that fold’s training. The final model was refitted on the complete training split (all 80% of clusters) after cross-validation.

### Performance Evaluation

Model performance was assessed on the held-out test set and independently on the C207 benchmark using five metrics. Hit Rate top 10% is the fraction of labeled sites recovered within the top 10% of ranked candidates per complex. Mean reciprocal rank (MRR) and the average and median rank of the labeled target residue capture how highly the true target was ranked. Search space reduction is the proportional decrease from the total nucleophile-capable residue count to the rank position of the true target. Normalized discounted cumulative gain (NDCG) captures overall ranking quality. All metrics were computed per query group (protein–warhead pair) and aggregated both overall and by target residue type, with out-of-fold predictions used throughout to avoid inflated in-sample estimates.

### Ablation Study

To assess the individual contribution of each physicochemical feature to ranking performance, we performed a leave-one-out ablation in which each feature group was removed in turn, and the model was retrained and re-evaluated under the same cross-validation and held-out test protocol. These features include solvent accessibility/desolvation cost, N-terminus status, residue identity, deprotonation state, and quantum reactivity descriptors (all six SQM features: nucleophilicity index, Fukui electrophilicity index, HOMO–LUMO gap, partial charge, electrophile LUMO energy, and nucleophile HOMO energy). Each ablation arm was evaluated against the full-feature model using identical cluster-level splits to isolate the effect of feature removal from variation in train/test split in terms of hit rate in the top 10% shortlist and normalized discounted cumulative gain at 15, NDCG@15, which measures ranking quality. Hit rate was evaluated using a percentile-based cutoff scaled to each complex’s candidate pool size, consistent with the varying size of the nucleophile-capable residue pool across structures. NDCG@15 was evaluated using a fixed top-15 cutoff, following conventional NDCG usage, and provides a complementary, pool-size-independent measure of ranking quality near the top of the list. Because each query group has exactly one labeled true target, NDCG was computed using binary relevance (the true target residue receives a relevance score of 1; all other candidates receive a score of 0).

### Model Validity and Feature Contributions

We examined the contribution of physicochemical features in *CovSite* to overall performance through a leave-one-out ablation, in which each feature group was removed in turn, and the model was retrained and re-evaluated under identical cluster-level cross-validation splits (**Figure 2A, B**). N-terminus status had the highest cost of any feature group by a wide margin, reducing the top 10% hit rate from 88.7% to 54.5% (Δ34.2 points) and NDCG@15 from 0.498 to 0.339 (Δ0.159) upon removal. This is significantly larger than the next most costly feature, solvent accessibility (SASA), which reduced hit rate to 69.7% (Δ19.0) and NDCG@15 to 0.380 (Δ0.118) when left out. Quantum reactivity descriptors showed a comparatively modest hit-rate cost (85.0%, Δ3.7) but a disproportionately larger NDCG cost (0.387, Δ0.111), nearly matching SASA’s NDCG reduction despite a hit-rate cost roughly one-fifth the size, indicating that QM reactivity descriptors primarily improve candidate ranking already in the shortlist (top 10%) rather than controlling which candidates make it in. Deprotonation probability and residue-type identity showed the smallest individual costs (hit rate Δ1.8 and Δ0.4, respectively; NDCG Δ0.027 and Δ0.010), consistent with these features contributing real but comparatively modest marginal signal once the remaining feature set is present. Deprotonation probability is estimated by a separately trained classifier conditioned on local pKa-relevant structural features rather than derived from the QM descriptors themselves, so the two feature groups are mechanistically independent. The combined removal of QM reactivity and deprotonation results in a lower NDCG (Δ0.077) than the removal of QM reactivity alone (Δ0.111); this is reported descriptively rather than interpreted as evidence of redundancy between the two feature groups.

**Figure 2.**
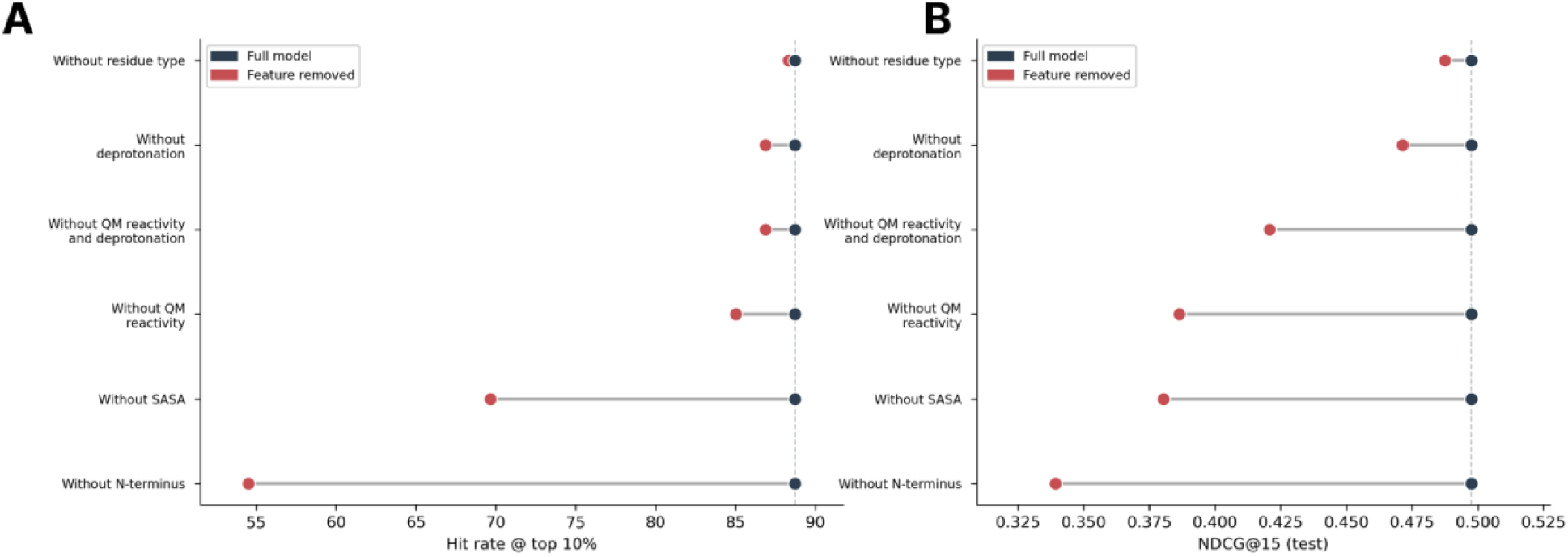
Leave-one-out feature ablation. (A) Hit rate at top 10% (B) NDCG@15 for full model (blue) versus full model with each feature group removed and retrained (red) under identical cluster-level cross-validation splits.

SHAP (SHapley Additive exPlanations), which quantifies the absolute strength of a feature’s impact on model predictions, was used to determine which variables have the greatest impact on model performance. The magnitude of the N-terminus ablation effect is notable given that N-terminus status ranks lowest by mean SHAP magnitude among all features^14^ (**Figure 3A**). This apparent discrepancy reflects the sparse, specific nature of N-terminus status, which is highly informative for a dominant subset of N-terminal threonine’s in Ntn-hydrolases, whose nucleophile activation proceeds via a free α-amino group rather than via conventional side-chain deprotonation. Mean SHAP magnitude reflects the average importance across the full candidate population. In contrast, leave-one-out ablation reflects cost concentrated in the small subgroup of residues (such as N-terminal threonine’s) where that feature integration results in a noticeable shift in accuracy. The two measures are expected to verge exactly in cases, such as these, where a feature’s relevance is concentrated in a narrow subset of residues rather than spread evenly across all of them. To rule out test-set-specific variation as a confound, we compared the held-out test set’s overall hit rate (88.7%) against the distribution of hit rates across the 5 cross-validation folds used during model development (**Figure 3B**). The held-out value fell within the interquartile range of the fold distribution and close to the fold median, indicating that test-set performance is consistent with expected cross-validation variance rather than reflecting an unusually favorable or unfavorable held-out split.

**Figure 3.**
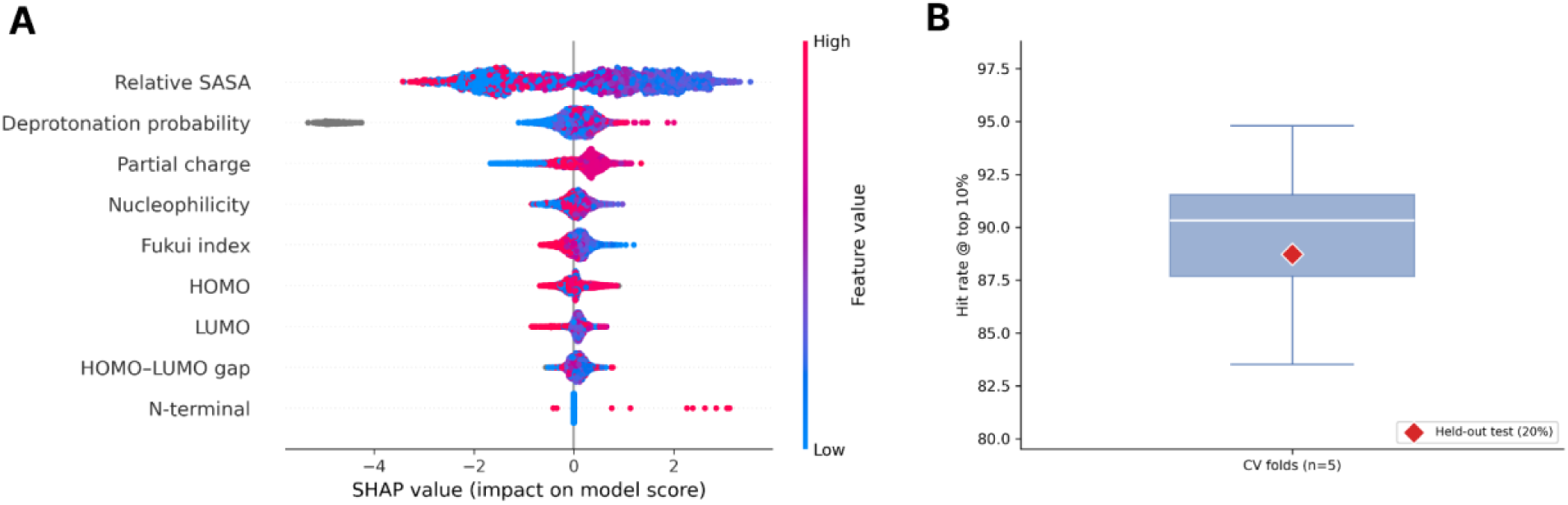
SHAP feature importance and cross-validation consistency. (A) SHAP summary plot showing distribution of SHAP values for each feature. (B) Hit rate at top 10% across 5 cross-validation folds used during development (box: IQR, whisker: range) versus 20% held-out clusters for testing.

### Residue-Type Specificity and Search-Space Composition

Because the model has access to residue identity as one of its features, we checked whether its predictions track the true target’s residue type, rather than defaulting to a few convenient residues (**Figure 4**). Looking at only query groups with confirmed residue-type labels, the model’s top pick matched the correct residue type in 88% of Cys-target proteins, 90% of Ser-target proteins, and 67% of Lys-target proteins (**Figure 4C**), all higher than would be expected by chance alone. We also measured how concentrated each residue type was among the top 10% of ranked candidates, relative to its overall frequency across all proteins. By this measure, the true target residue type was over-represented in the shortlist by 5.02-fold for Cys, 2.02-fold for His, 1.83-fold for Lys, and 2.74-fold for Ser (**Figure 4B**, diagonal).

**Figure 4.**
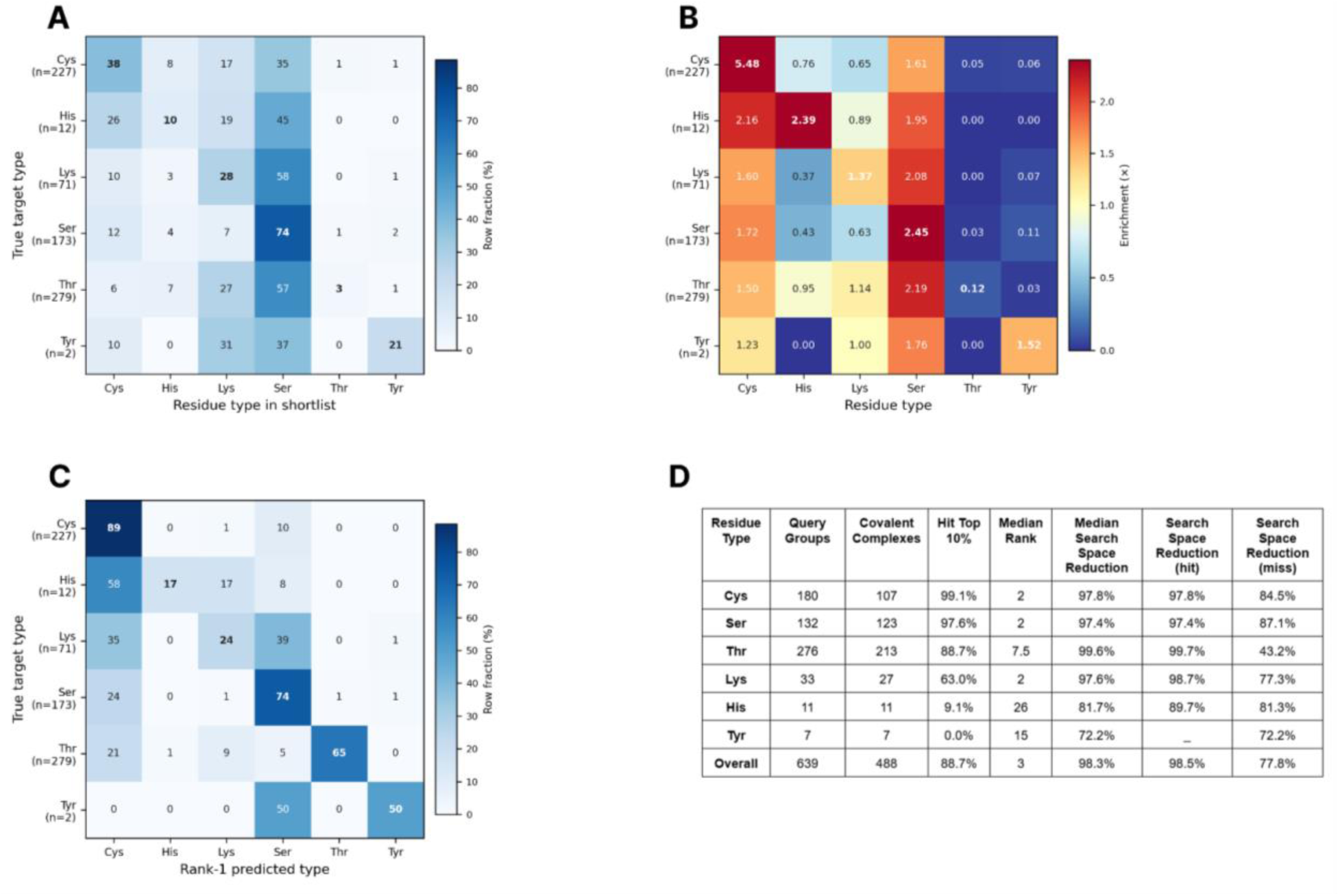
Residue-type specificity of CovSite on held-out test set. (A) Composition of the top 10% shortlist by residue type (columns) compared by true target residue type (rows). (B) Enrichment of each residue type within top 10% shortlist relative to baseline frequency among all residues for that protein. (C) Confusion matrix of rank 1/top-ranked predicted residue type versus true target residue type. (D) Per-residue-type test performance (note: Query Groups are warhead-residue pairs).

Thr and Tyr broke from this pattern. Thr showed almost no enrichment across the entire residue-type column, including for its own true targets (0.12-fold), and the model’s top pick matched the true Thr target only 55% of the time (**Figure 4C**), most often confusing it with Lys (21%) or Cys (15%). This matches what we see in the per-residue accuracy results (**Figure 4D**), where Thr had the largest gap between the overall hit rate (88.7%) and the extent of the misses. The likely explanation is that threonine reactivity is uneven as a class. Most candidate threonines in a given protein are ordinary, non-reactive residues, whereas reactive threonine instances are concentrated almost entirely in a rare subgroup: N-terminal threonines. Seen this way, the lack of enrichment for Thr is not a sign that the model is failing on threonine, but instead a sign that the model correctly avoids favoring threonine candidates indiscriminately, since most are genuinely non-reactive. Tyr’s near-zero enrichment (0.06-fold) and 0% top-pick accuracy are best explained by the limited Tyr data available for evaluation (n=7 query groups), so these results should be treated as inconclusive given the sample size rather than as a reliable estimate of how the model handles tyrosine.

Per-residue test accuracy (**Figure 4D**) showed hit rates of 99.1% (Cys, n=180), 97.6% (Ser, n=132), 88.7% (Thr, n=276), 63.0% (Lys, n=33), 9.1% (His, n=11), and 0.0% (Tyr, n=7), with the candidate pool shrinking by more than 97% for Cys, Ser, and Thr. Search-space reduction conditioned on outcome diverged substantially between hits and misses, particularly for Thr (97.8% reduction when hit vs. 43.2% when missed) and Lys (98.7% vs. 77.3%), indicating that when the model fails for these residue types, the true target tends to fall considerably further down the ranked list than for Cys or Ser failures, rather than narrowly missing the top-10% cutoff.

### Independent Benchmark Evaluation

To evaluate *CovSite* against existing covalent screening methods on a common reference, we assessed performance on the C207 benchmark set, an independent collection of 207 cysteine-targeted covalent complexes from Scarpino et al. with no protein-ID overlap with the *CovalentInDB 2.0* training and cross-validation sets (**Figure 5**). Because C207 labels are exclusively cysteine, which reflects the residue scope of the docking tools it was originally curated to benchmark, comparison on this set evaluates *CovSite* specifically on its cysteine-targeting performance. However, candidate pools and rankings were not restricted to cysteine, and *CovSite’s* predictions were generated by ranking the true cysteine target against all co-occurring nucleophile-capable residues (Cys, Ser, Thr, Tyr, Lys, His) on each structure, consistent with its fully blind operating mode.

**Figure 5.**
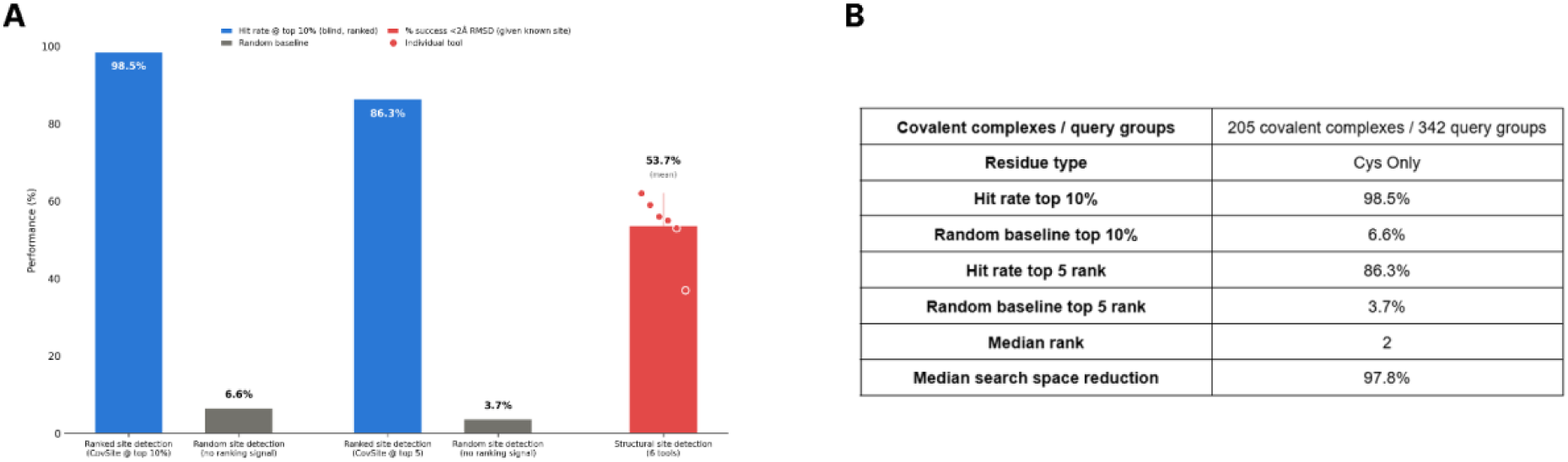
Comparison of method accuracy on hit detection with blind ranking versus structural docking on C207 benchmark. (A) Performance on 205 labeled cysteine covalent sites from C207 benchmark. Blue bars: CovSite’s hit rate at top 10% (98.5%) and top 5 rank (86.3%) candidates. Grey bars: per-site random chance baseline. Red bar: mean of best-scoring poses at ≤2 Å RMSD (53.7%) across six structure-based screeners (ICM-Pro, CovDock, FITTED, AutoDock4, GOLD, MOE). (B) Summary of CovSite evaluation on C207.

*CovSite* correctly identified the confirmed covalent site within the top 10% of ranked candidates in 98.5% of the 205 labeled sites, and within a fixed top 5 rank cutoff in 86.3% of cases (**Figure 5A**). The true target rank had a median score rank of 2 and a strong median search space reduction of 97.8% of the six nucleophilic residues (**Figure 5B**). Some labeled sites are targeted by more than one warhead chemotype, and each chemotype was evaluated as its own separate query. A site was counted as successfully recovered only if *CovSite* correctly identified all its associated warhead-specific query groups, not just one. This stricter, “all-warheads-must-match” methodology means the odds of recovering a site by pure chance are lower for sites with multiple warheads than for sites with just one. To account for this, the chance baseline was calculated separately for each site as the product of the random-hit probabilities for its individual query groups, rather than using a single fixed percentage across all sites. The resulting baseline was 6.6% at top 10% and 3.7% at top 5 rank (**Figure 5A**), far below *CovSite’s* observed performance at both thresholds and confirming that recovery is not attributable to chance.

On the same C207 set, six structure-based covalent screening tools achieved a mean best-scoring pose success of 53.7% at ≤2 Å RMSD, ranging from 37% to 62%. CovSite and these structural tools demonstrate two distinct methods for hit-detection of a ligand on a protein’s target site. CovSite utilizes blind residue detection with ligand warhead chemical reactivity and structural properties, whereas docking tools assess whether a specified electrophile can achieve a geometrically viable pose on a known target site for a covalent bond.

### Recapitulating Known Covalent Binding Sites

To assess whether *CovSite*’s blind ranking recapitulates experimentally established covalent binding sites across mechanistically distinct reactivity classes, we examined two case studies spanning a canonical cysteine side-chain target and an N-terminal, non-side-chain-mediated target. The two case studies presented here span the two mechanistically distinct reactive-site classes *CovSite* is designed to resolve: conventional side-chain nucleophilicity, exemplified by the disease-associated Cys12 in KRAS G12C, and N-terminal amine-mediated nucleophilicity, exemplified by the catalytic Thr1 of the proteasome β5 subunit. For the proteasome drug *carfilzomib*, *CovSite* correctly identified Thr1 (chain Z) as the most reactive site, ranking it first among 6,440 candidate residues in the protein. For the KRAS drug *sotorasib*, CovSite did the same for Cys12 (chain A), ranking it first out of 46 candidate residues. In both cases, *CovSite* identified the one true binding site from hundreds of possibilities with no prior knowledge of where to look.

To test whether *CovSite* can correctly identify the residue target of an established protein/inhibitor pair under fully blind conditions, we screened the 20S proteasome using the carfilzomib and ixazomib warheads. The human constitutive 20S proteasome maintains cellular homeostasis by degrading intracellular proteins, including those involved in cell cycle regulation and apoptosis. This has led to the 20S proteasome becoming an oncology target; covalent proteasome inhibitors inactivate β5 subunit active sites, disrupting cell cycle control to the detriment of actively proliferating cancer cells. Mechanistically, the proteasome β5 subunit is covalently inhibited by FDA-approved drugs Carfilzomib and Ixazomib through nucleophilic attack of the catalytic Thr1 free α-amino group by the drug’s warhead, which are epoxyketone and boronic acid, respectively. *CovSite rank*ed the experimentally validated nucleophilic residue among the top 10 residue targets for each drug, ranking it 6th for Carfilzomib and 1st for Ixazomib (**Figure 6**). The 5 residues that outscored Thr1 for Carfilzomib were each N-terminal threonine over other chains. Favorable scoring in this evaluation shows that *CovSite* can correctly prioritize an N-terminal threonine over higher-occupancy or more solvent-exposed cysteine and serine residues elsewhere on the subunit. Evaluation on the 20s proteasome demonstrates successful use of *CovSite* on a very large, multi-subunit structure.

**Figure 6.**
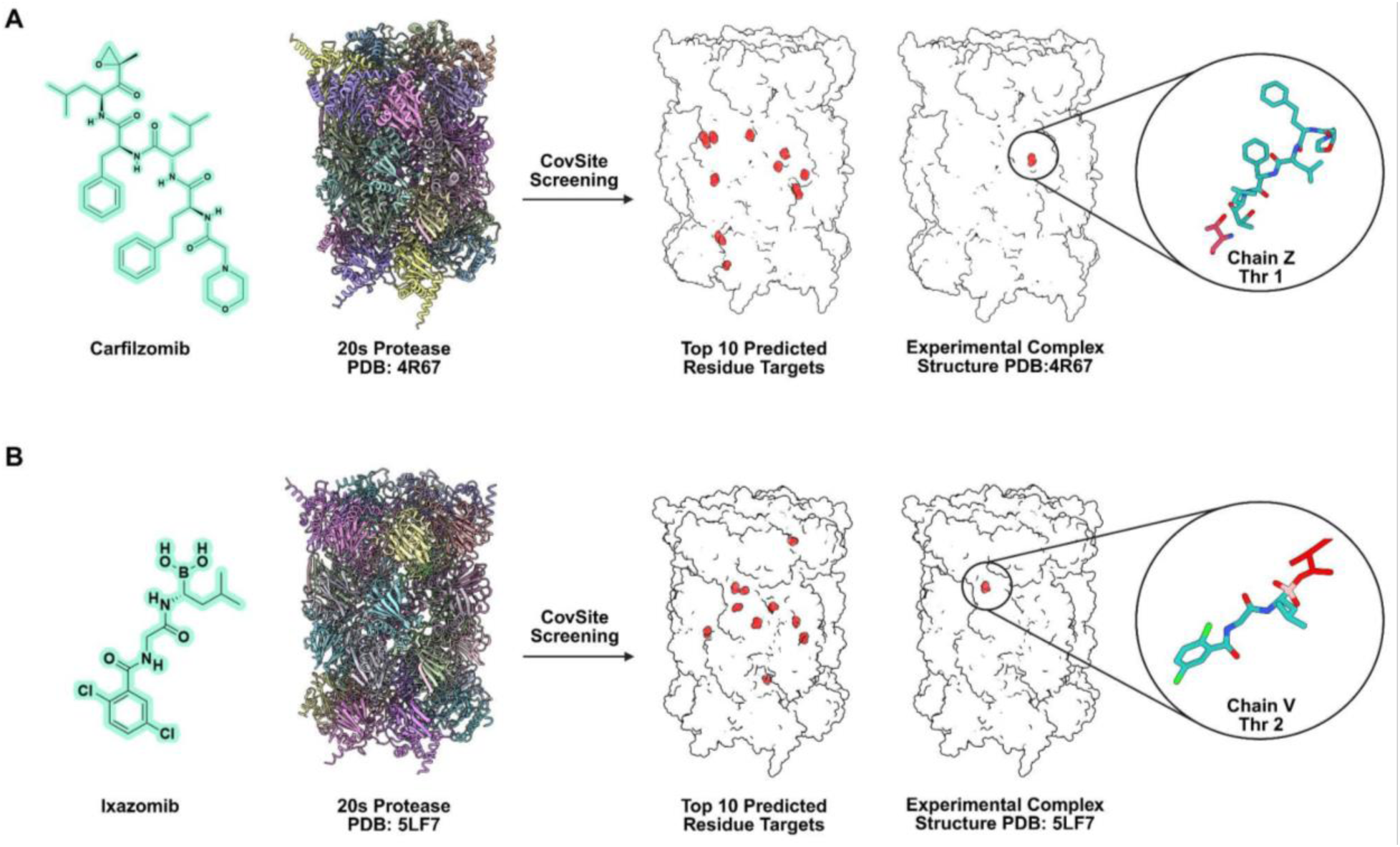
*CovSite* was run in fully blind mode against the unliganded target structure, with no prior specification of binding site, residue identity, or warhead chemistry, and the rank assigned to the known clinical target residue (Thr1 for the proteasome β5 subunit) was recorded among all nucleophile-capable candidates on the structure. Tested with two known covalent inhibitors of Carfilzomib (A) and Ixazomib (B).

Further, we expanded the scope of the search space to test whether *CovSite* can identify nucleophilic residues in proteins that closely resemble a structure in the training set; we screened K-Ras G12C with the Sotorasib warhead after training CovSite on H-Ras, which has 95% sequence similarity to K-Ras. K-Ras, an oncogenic protein, is known as “the most mutated protein in cancer,” and mutagenesis of the K-Ras isoform is the most prevalent of Ras mutations, of which roughly 30% of all human cancers harbor. The K-Ras G12C mutant is covalently inhibited by Sotorasib through Michael addition of the Cys12 thiolate to the drug’s acrylamide warhead, positioned within the switch II pocket adjacent to the nucleotide-binding site, exploiting a cysteine introduced by the G12C oncogenic mutation that is absent in both H-Ras and wild-type K-RAS. Even after being trained on a covalently bound H-Ras structure, CovSite correctly scored C12 as the top residue for the K-Ras G12C mutant **(Figure 7)**. This test demonstrates how CovSite can be used to screen residues of disease-associated mutations for potential covalent inhibition, even when the dominant reactive site contains multiple surface-exposed cysteines. Furthermore, this demonstrates the ability to detect a mutated site in an oncogenic protein that a current covalent drug (Sotorasib) can target. Both case studies recapitulated the experimentally confirmed site within the top ten candidates, demonstrating that blind ranking performance generalizes across mechanistically distinct reactivity classes (sidechain vs. N-terminal amine-mediated nucleophilicity) and across disease contexts (oncology vs. proteasome inhibition).

**Figure 7.**
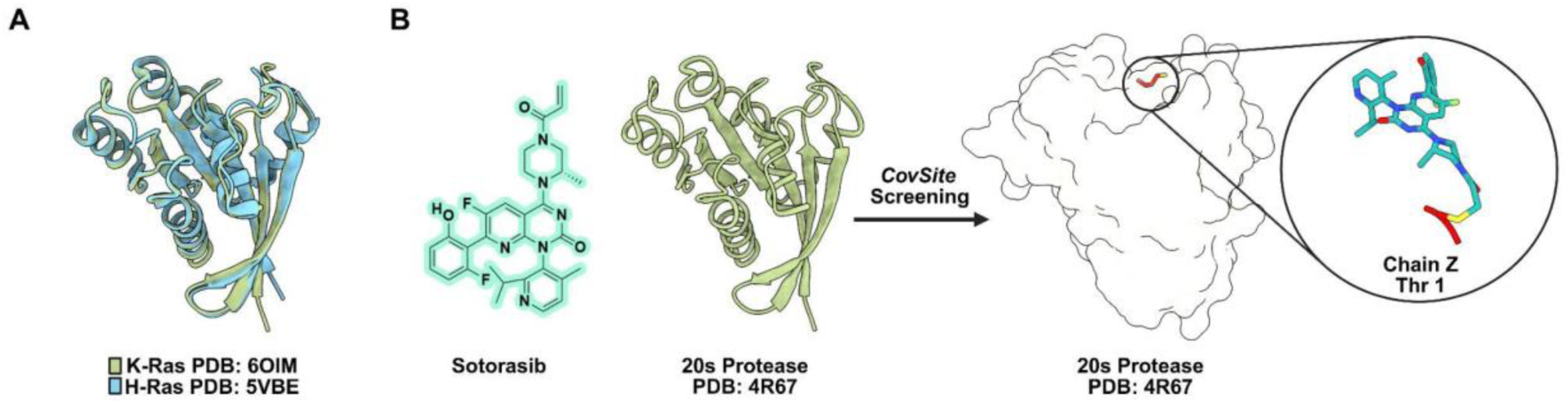
*CovSite* was run in fully blind mode against the target structure (A) with the ligand (B), with no prior specification of binding site, residue identity, or warhead chemistry, and the rank assigned to the known clinical target residue (Cys12 for K-Ras G12C) was recorded among all nucleophile-capable candidates on the structure.

## Discussion

*CovSite* was developed and evaluated using protein-cluster-grouped cross-validation (MMseqs2, 50% sequence identity) to prevent data leakage between training and evaluation. The model achieved an 88.7% hit rate at top-10% on the held-out test set, with held-out performance falling within the interquartile range of the 5-fold cross-validation distribution, indicating that test-set results reflect expected model behavior rather than a favorable held-out split (**Figure 3A**, **Figure 4D**). Leave-one-out ablation showed that removing N-terminus status or solvent accessibility each caused a substantial, independent reduction in performance, and quantum reactivity descriptors contributed meaningfully to NDCG (ranking quality) even where their contribution to accuracy (hit rate) was comparatively modest. Quantum reactivity descriptors better rank the candidates already filtered through previous stages rather than determine which candidates make the 10% top-hit list.

External validity was assessed using the independent C207 benchmark, curated to have no protein-identifier overlap with the CovalentInDB 2.0 development set^2^. Six tools (AutoDock4, CovDock, FITTED, GOLD, ICM-Pro, MOE) were each provided with the target binding site in advance and evaluated on whether their top-ranked pose reproduced the crystallographic binding geometry within 2 Å RMSD. On the other hand, *CovSite* narrows down residues without structural docking, using site-level chemical reactivity and accessibility. These two distinct methods were compared for the use case of hit detection when exploring potential ligands (**Figure 5A).** Strong results of 98.5% recapitulation at the top 10% of residues, with a median rank of 2 for the target site on C207 (**Figure 5A**, **Figure 5B**), demonstrate that residue detection using a non-structural, machine-learning model can be an effective method for identifying viable warheads on target residues. Furthermore, both case studies involving covalent drugs and clinically relevant proteins highlight the capability that *CovSite* extends across diverse disease contexts and drug-reaction mechanisms (**Figure 6**, **Figure 7**).

A comparison of these two methods was used because existing covalent docking tools are the closest available point of comparison for site-level performance in this field, and no benchmark or tool directly mirrors the blind, warhead-specific screening task of *CovSite*. This lack of benchmarks further highlights a broader gap in covalent drug discovery: determining whether a given warhead is chemically and structurally compatible with the target site before incurring the costs of more expensive structural modeling or experimental validation. Evaluation of warhead reactivity towards the target site can facilitate the most popular drug discovery approach of attaching an electrophilic warhead to known, high-affinity non-covalent binders^10^. The limitation of these findings, however, is the absence of decoy warheads to serve as negative controls for these active ligands, due to the lack of available covalent datasets. Although recent resources (e.g., COValid) have expanded the set of covalent benchmarks, current datasets still have a limited number of warhead chemotypes and are dominated by cysteine-reactive compounds. While Cys-reactive nucleophilic residues are useful for site-specific chemistry, such as SnapTag for protein-protein assemblies^15,16^ and KRAS^G12C^ for cancer immunotherapy^3,17^, exploring novel protein-based inhibitors beyond this residue remains a challenge in the field. In the case of COValid, warhead diversity was purposely constrained to be one chemotype because existing screening tools generally do not account for covalent reactivity, making ranking multi-warhead libraries infeasibl^18^.

To help mitigate this limitation, *CovSite* ranks across all six canonical nucleophile-capable residue types (here, Cys, Ser, Thr, Tyr, Lys, His), rather than restricting predictions to solely cysteine. This broadens the diversity of protein-electrophile interactions evaluated and tests plausible reactive sites beyond a single residue, introducing a greater set of negative residue-warhead comparisons than a cysteine-only search. Despite this increased search space, *CovSite* maintains target residue-type specificity, as shown by the residue distribution in **Figure 4A-C**. Furthermore, covalent reactivity is not confined to cysteine even though cysteine-targeting chemistry dominates current clinical and tool-development efforts. Performance was strongest for cysteine and serine, consistent with their greater representation in the training distribution, and rank-1 predictions tracked the true target’s residue type well above chance for cysteine, serine, threonine, and lysine, indicating that the *CovSite* residue-type-aware predictions reflect target-specific signal rather than defaulting to any convenient residue types. As coverage of data on non-cysteine covalent targets in the literature expands, retraining on larger, more balanced datasets is expected to improve performance for underrepresented residue types, particularly histidine and tyrosine, for which current results should be treated as preliminary. Additionally, the physicochemical descriptors in *CovSite* capture side-chain-level and chain-positional reactivity context, but activation mechanisms outside this scope, such as unanticipated cofactor- or metal-assisted activation, would not be captured by the current feature set without targeted feature engineering, as was done for the N-terminal case.

Lastly, another limitation is that *CovSite* is not a structural or pose-prediction model. It does not generate binding poses, predict interaction geometry, or provide distance and angle constraints between the electrophile and the residue, and applications that require this information would need downstream covalent docking. An effective workflow could use *CovSite* as an upstream warhead-identification step before testing the geometric feasibility with current covalent docking tools, expanding the discovery of covalent inhibitors and protein-inhibitor pairs for applications in cell engineering^19^, drug discovery^20^, and bioconjugation^21,22^. In the current form of our platform, this is an intentional scope boundary, where pose-level structural modeling carries computational costs that would compromise the high-throughput, blind-screening role *CovSite* is designed to fill^5^.

## Conclusion

*CovSite* addresses a gap in covalent drug discovery that conventional pocket-based virtual screening and existing covalent docking tools are not built to fill: blind, residue-level identification of covalent reactivity toward a specified electrophile, without prior knowledge of the binding site. Using a single learned ranking model over physicochemical descriptors spanning solvent accessibility, protonation state, quantum reactivity, and structural context, *CovSite* achieved an 88.7% hit rate at top 10% on held-out cross-validation data (**Figure 3B)** and generalized to an independent benchmark, recovering 98.5% of confirmed covalent sites on the C207 set while reducing the candidate search space by a median of 97.8% (**Figure 5A, B**). Existing covalent docking tools, evaluated on the same benchmark under the more favorable condition of a pre-specified binding site, achieved best scoring pose accuracies of 53-62%, addressing a distinct but related condition for covalent inhibitor viability. Beyond cysteine, *CovSite* extends ranking coverage to five additional canonical nucleophilic residue types, addressing a systematic gap in a field where high-throughput tools remain largely cysteine-specific. Ablation and residue-specificity analyses confirm that this coverage reflects a genuine, target-aware reactivity signal rather than reliance on residue identity alone. As covalent structural databases expand to better represent underrepresented nucleophiles and reactivity descriptors continue to improve, the single-model architecture offered by *CovSite* enables targeted refinement without restructuring the framework, positioning it as a practical first-pass screening tool for early-stage covalent drug discovery, particularly for novel or understudied targets with limited binding-site information.

## AUTHOR INFORMATION

### Corresponding Author

Blaise R. Kimmel, Ph.D. 151 W. Woodruff Avenue Columbus, OH 43210

### Author Contributions

The manuscript was written through the contributions of all authors. All authors have given approval to the final version of the manuscript. B.R.K. supervised the project and acquired funding to support the research.

## Acknowledgements

This work was supported in part by The Ohio State University Center for Cancer Engineering, Curing Cancer Through Research in Engineering and Sciences, and the National Cancer Institute (NCI) through the R21 grant 1R21CA312456. B.R.K. acknowledges financial support from the Prostate Cancer Foundation Young Investigator Award. This work used the Ohio Supercomputer Center, which provides High Performance Computing resources and expertise to academic researchers across the State of Ohio. OSC is a member of the Ohio Technology Consortium, a division of the Ohio Department of Higher Education.

## Code Availability

https://github.com/Kimmel-Lab/CovSite

## References

1. Singh, J., Petter, R. C., Baillie, T. A. & Whitty, A. The resurgence of covalent drugs. Nat. Rev. Drug Discov. 10, 307–317 (2011).

2. Du, H. et al. CovalentInDB 2.0: an updated comprehensive database for structure-based and ligand-based covalent inhibitor design and screening. Nucleic Acids Res. 53, D1322–D1327 (2025).

3. Chen, H. et al. A Perspective on Covalent Inhibitors: Research and Development Trends of Warheads and Targets. Preprint at 10.26434/chemrxiv-2023-wf79n (2023).

4. Ghosh, A. K., Samanta, I., Mondal, A. & Liu, W. R. Covalent Inhibition in Drug Discovery. ChemMedChem 14, 889–906 (2019).

5. Singh, N., Vayer, P. & Villoutreix, B. O. The covalent docking software landscape: features and applications in drug design. Brief. Bioinform. 26, bbaf697 (2025).

6. Lionta, E., Spyrou, G., Vassilatis, D. & Cournia, Z. Structure-Based Virtual Screening for Drug Discovery: Principles, Applications and Recent Advances. Curr. Top. Med. Chem. 14, 1923–1938 (2014).

7. Skolnick, J., Gao, M., Roy, A., Srinivasan, B. & Zhou, H. Implications of the small number of distinct ligand binding pockets in proteins for drug discovery, evolution and biochemical function. Bioorg. Med. Chem. Lett. 25, 1163–1170 (2015).

8. Sotriffer, C. Docking of Covalent Ligands: Challenges and Approaches. Mol. Inform. 37, 1800062 (2018).

9. Oyedele, A.-Q. K. et al. Docking covalent targets for drug discovery: stimulating the computer-aided drug design community of possible pitfalls and erroneous practices. Mol. Divers. 27, 1879–1903 (2023).

10. Keeley, A., Petri, L., Ábrányi-Balogh, P. & Keserű, G. M. Covalent fragment libraries in drug discovery. Drug Discov. Today 25, 983–996 (2020).

11. Scarpino, A., Ferenczy, G. G. & Keserű, G. M. Comparative Evaluation of Covalent Docking Tools. J. Chem. Inf. Model. 58, 1441–1458 (2018).

12. Dampalla, C. S. et al. Structure-Guided Design of Potent Inhibitors of SARS-CoV-2 3CL Protease: Structural, Biochemical, and Cell-Based Studies. J. Med. Chem. 64, 17846–17865 (2021).

13. Hasan, M. N., Ray, M. & Saha, A. Landscape of *In Silico* Tools for Modeling Covalent Modification of Proteins: A Review on Computational Covalent Drug Discovery. J. Phys. Chem. B 127, 9663–9684 (2023).

14. NIPS-2017-a-unified-approach-to-interpreting-model-predictions-Paper.

15. Metcalf KJ et al. Synthetic Tuning of Domain Stoichiometry in Nanobody-Enzyme Megamolecules. Bioconjug. Chem. 32, 143–152 (2021).

16. Vu, T. Q. et al. Controlled Assembly of Vesicle-Based Superstructures Using Megamolecules. ACS Appl. Mater. Interfaces 18, 4937–4951 (2026).

17. Huang, F., Han, X., Xiao, X. & Zhou, J. Covalent Warheads Targeting Cysteine Residue: The Promising Approach in Drug Development. Molecules 27, 7728 (2022).

18. Shamir, Y. et al. Discovery of Covalent Ligands with AlphaFold3. J. Am. Chem. Soc. 148, 13043–13054 (2026).

19. Galvan, S., Teixeira, A. P. & Fussenegger, M. Enhancing cell-based therapies with synthetic gene circuits responsive to molecular stimuli. Biotechnol. Bioeng. 121, 2987–3000 (2024).

20. Boike, L., Henning, N. J. & Nomura, D. K. Advances in covalent drug discovery. Nat. Rev. Drug Discov. 21, 881–898 (2022).

21. Chen, Y. et al. Molecular Matchmakers: Bioconjugation Techniques Enhance Prodrug Potency for Immunotherapy. Mol. Pharm. 22, 58–80 (2025).

22. Kimmel, B. R. & Mrksich, M. Development of an Enzyme-Inhibitor Reaction Using Cellular Retinoic Acid Binding Protein II for One-Pot Megamolecule Assembly. Chem. – Eur. J. 27, 17843–17848 (2021).

