## Supplementary material for "*CovSite*: A High-Throughput Blind Covalent Screening Framework for Reactive Site Detection": SI

**Supplementary Information**

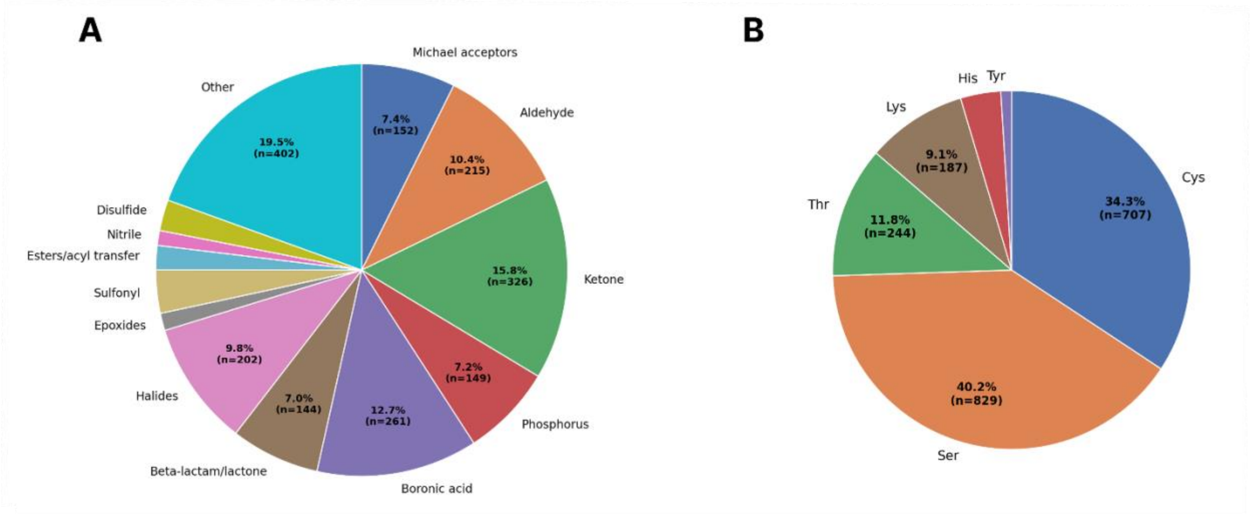


**Supplementary Figure 1. Composition of curated CovalentInDB 2.0 training set.** (A) Distribution of warhead chemotypes among 2,062 curated entries. (B) Distribution of the six nucleophilic residues of interest: Cys, Ser, Thr, Lys, His, Tyr.

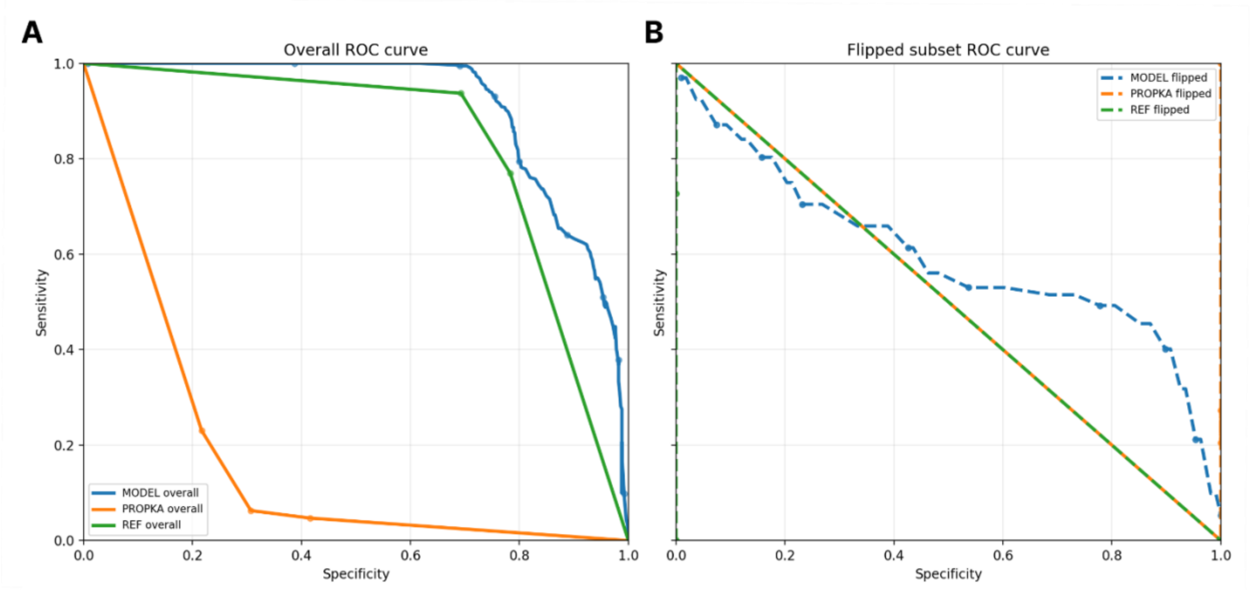


**Supplementary Figure 2. The model retains sensitivity to environment-driven protonation shifts, compared to PROPKA.** ROC curves (sensitivity vs. specificity), swept across pH thresholds for trained deprotonation model (MODEL), PROPKA, and the reference pKA baseline (REF), evaluated on the PKAD-R dataset. (A) Overall ROC curve across the six nucleophilic residues of interest. (B) ROC curve restricted to the flipped subset, where environmental context changes the residue’s protonation state relative to its pKa baseline. PROPKA and REF are indistinguishable, consistent with PROPKA failing to capture environment-driven shifts in protonation.
